# Metagenome-based diversity and functional analysis of culturable microbes in sugarcane

**DOI:** 10.1101/2024.08.05.606635

**Authors:** Haidong Lin, Liang Wu, Lijun zhang, Ta Quang Kiet, Peng Liu, Jinkang Song, Xiping Yang

## Abstract

Sugarcane, widely acknowledged as the foremost crop for sugar and energy production on a worldwide scale, is confronted with many diseases that pose serious threats to its production. Biological control has become more popular as an approach for preventing and controlling diseases because of its environment-friendly characteristics. However, there is a lack of thorough investigation and use of microbial resources in sugarcane. This study conducted a thorough analysis of culturable microbes and their functional features in different tissues and rhizosphere soil of four diverse sugarcane species using metagenomics techniques. The results revealed significant microbial diversity in sugarcane’s tissues and rhizosphere soil, including several important biomarker bacterial taxa identified, which are reported to engage in several processes that support plant growth. The LEfSe studies identified unique microbial communities in different parts of the same sugarcane species, particularly *Burkholderia*, which exhibited significant variations across the sugarcane species. Microbial analysis of carbohydrate-active enzymes (CAZymes) indicated that genes related to sucrose metabolism were mostly present in specific bacterial taxa, including *Burkholderia, Pseudomonas, Paraburkholderia*, and *Chryseobacterium*. This study improves understanding of the diversities and functions of endophytes and rhizosphere soil microbes in sugarcane. Moreover, the approaches and findings of this study provide valuable insights for microbiome research and the use of comparable technologies in other agricultural fields.

## Introduction

Sugarcane (*Saccharum* spp.) is a critical global sugar and energy crop, and its yield and quality significantly impact the global economy and food industry. However, sugarcane production is often endangered by a wide range of diseases, and the prevention and control of sugarcane diseases is an urgent issue for the industry. Thus, developing effective disease control strategies for sugarcane diseases is particularly important. Chemical pesticide control in traditional sugarcane agriculture can increase yields temporarily, but it will lead to issues like pathogen resistance, excessive pesticide residues, and environmental contamination. Biological control, as an environmentally friendly alternative, has gradually become one of the popular research directions in recent years. Because of the potential benefits to the environment and human health, the use of plant endophytes and rhizosphere microbes for disease prevention has gained a lot of attention. Consequently, utilizing rhizosphere microbial resources and plant endophytes is essential for disease prevention.

Plant endophytes and rhizosphere microbes are two key groups of sugarcane that play an extremely important role in promoting sugarcane growth and improving stress tolerance (1, 2). These microbes play multiple roles in the growth of sugarcane, such as facilitating nutrient uptake (3), production of growth regulators (4), inhibition of pathogenic microbes (5), and enhancement of tolerance to abiotic stresses (6), etc. The main growth-promoting microbes of sugarcane identified so far include *Rhizobium, Burkholderia, Streptomyces, Enterobacter, Pseudomonas*, and *Bacillus* (7, 8). Understanding the diversity of these microbes and their functions is essential for green production of sugarcane.

Metagenomics is a method that allows for the comprehensive analysis of all microbial genomes in environmental samples (9). It provides a means of systematically unravelling microbial species’ diversity, functional attributes, and interrelationships, thereby deepening the understanding of their functions in ecosystems and evolutionary processes (10). Despite significant advances in other areas of metagenomics, such as the human microbiome (11), soil microbiome (12), and marine microbiome (13), its application in sugarcane microbiome research is still limited. Current researchers have initiated investigations into the makeup of the sugarcane microbiome. However, most of these studies primarily concentrate on specific microbial taxa or functions, and there is a paucity of comprehensive metagenomics studies. This current study aims to comprehensively investigate the diversity and roles of endophytes and rhizosphere soil microbes in sugarcane, and furthermore, to discover crucial functional microbes and biocontrol factors that can be used to develop novel microbial resources for the biocontrol of sugarcane diseases and the enhancement of sugarcane growth.

With the rapid development of bioinformatics and biotechnology, metagenomics-based research will provide powerful support for the sustainable production of sugarcane and even other crops. The innovation of this study is the application of metagenomics technology to the endophytic and rhizosphere soil culturable microbes of different species of sugarcane for microbial diversity and related functional analyses, which can help to reveal the microbial resources with potential application value more precisely. In this study, 48 culturable microbial samples obtained from different parts of four sugarcane species (*S. spontaneum, S. robustum, S. officinarum*, and *S*. hybrid) were isolated and subjected to metagenome sequencing, and significantly different microbes were identified by comparing the differences in microbial composition and function under different environmental conditions. The study findings will not only offer novel approaches and techniques for the biological management of sugarcane, but also establish a scientific foundation and offer technological assistance for the advancement of microbial fertilizers and biopesticides.

## Materials and methods

### Sample collection and pre-processing

In order to ensure the representativeness of the samples, four species of the sugarcane genus (*S. spontaneum, S. robustum, S. officinarum* and *S*. hybrid) were selected for the experiment. Three genotypes were selected for each species, with a total of 12 samples, which were collected from the Guangxi Subtropical Agricultural Science New Base (Fusui County, Guangxi, China). In this study, the plant materials of sugarcane collected include leaves (the first leaf of the sugarcane plant that is fully green from the bottom to the top), stems (taken from the second node above the ground), and root samples (the soil still adhering to the roots was collected as rhizosphere soil samples by vigorous shaking). These samples were quickly placed in sterile bags after collection and stored in a 4°C incubator to preserve the microbes of the samples. They were then collected and processed within 24 hours.

### Isolation and culture of microbes

Four species of sugarcane materials were treated for culturable microbial isolation following the previous method (14). The isolated sugarcane culturable microbes (including endophytic microbes of stems, leaves, roots and rhizosphere soil microbes of the above four species of sugarcane materials) were aspirated and spread separately on five solid media (Nutrient Agar, Ashby’s Medium, Burke’s Medium, R2A Medium, Potato Dextrose Aga). The plates were inverted and incubated at 28°C for 72 h to facilitate colony growth.

### Metagenome DNA extraction and sequencing

A total of 48 microbial washed samples (sterile water was aspirated as a rinse solution using a sterile pipette in a horizontal flow clean bench) from different compartments and genotypes were obtained by collecting colonies after 72h incubation. The CTAB method was used to extract the total DNA of the sample genome, and DNA concentration and purity were tested on 1% agarose gel. Capillary electrophoresis was carried out using an AATI fragment analyzer to check the integrity of DNA. DNA was accurately quantified, and purity was determined using a Qubit fluorometer and Nanodrop kit to ensure the DNA samples suitable for subsequent high-throughput sequencing analysis library construction and sequencing. To construct Illumina sequencing libraries, we combined the NEBNext® UltraTM DNA Library Preparation Kit (New England Biolabs, USA) with 1 μg of DNA with index codes attached to the sequencing primers. The isolated DNA was sonicated and the ends were repaired to form a 350 bp fragment. After repairing the ends, adenine was added to the 3’ end of the DNA fragment, and then the adaptor sequence was ligated to both ends of the A-tailed DNA. The libraries were cleaned using AMPure XP technology from Beckman Coulter in Brea, CA. The size distribution and quantity of purified products were examined using an Agilent 2100 Bioanalyzer and real-time fluorescence quantitative PCR. After confirming library quality, all samples were subjected to paired-end sequencing using the Illumina NovaSeq 6000 platform with a read length of 150 base pairs (PE150).

### Data processing and analysis

The fastp tool (version: v0.19.4) was used to process the raw sequencing data to exclude low-quality base pairs, short sequences, and splice sequences (15). The comprehensive database Kraken2 (v2.1.1) was used to annotate the sample microbes by species. Bracken (v2.6.2) was then used to process the Kraken2 results and re-estimate abundance to corresponding species information and species-based abundance distributions, which formed the foundation for the diversity analysis (16, 17). To evaluate the diversity of the microbial community within a single sample, alpha diversity indices were computed, including the Shannon, Simpson, and Chao1 indices (18). The microbial community variations between samples were measured using beta diversity indices (19). Non-redundant genes were annotated by dbCAN and HMMER (v3.3.1) based on the CAZy database. The species composition and community results of grouped samples were examined for differences and relationships using PERMANOVA, the LEfSe statistical analysis approach, and other techniques.

## Results

### Diversity analysis of culturable microbes in sugarcane

For microbial isolation and cultivation, we chose four sugarcane species, with three different genotypes for each species. These isolated microbes included endophytes in the leaves, stems, roots and microbes in sugarcane rhizosphere soil. A total of 48 microbial samples were collected for metagenome sequencing (14). Using the α-diversity index, we performed a diversity study of culturable microbes in sugarcane based on the metagenome data. The rhizosphere soil microbes of the four sugarcane germplasm examined in this study exhibited a greater microbial diversity compared to other parts (Figure 1), suggesting the presence of a varied and abundant microbial community with a more uniformly distributed relative abundance. Among the endophytes of sugarcane, Simpson’s index showed a higher microbial diversity of culturable bacteria in the roots of *S. robustum* and in the leaves of *S. officinarum, S*. hybrid, and *S. spontaneum* (Figure 1B). Significant differences were observed in the Chao1 index of the microbial community between the various parts of *S. robustum* and *S. spontaneum* (*P* < 0.05) (Figure 1C). This phenomenon was especially evident in the stems and leaves of *S. robustum*, where the microbe abundance was significantly lower. The leaf endophytes of *S. robustum* exhibited a low diversity of microbial communities, as indicated by the low values of the Shannon, Simpson, and Chao1 indices, suggesting that specific microbial species were prominent, potentially influencing the plant’s development and physiological function.

**Figure 1:**
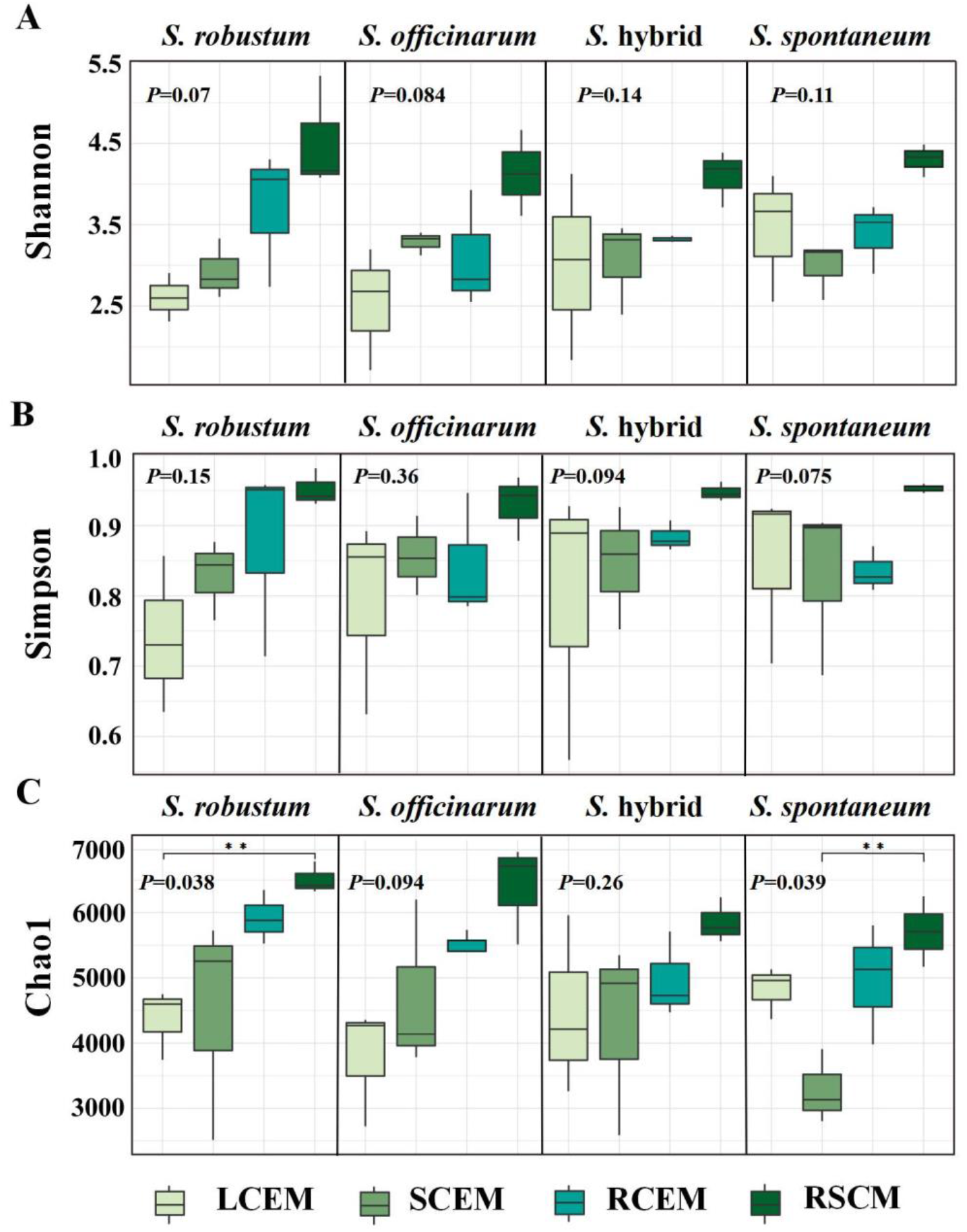
Shannon (A), Simpson (B), and Chao1(C) index of culturable microbial communities in different compartments of sugarcane (*S. spontaneum, S. robustum, S. officinarum, S*. hybrid) including LCEM (Leaf culturable endophytic microbes); SCEM (Stem culturable endophytic microbes); RCEM (Root culturable endophytic microbes). RSCM (Rhizosphere soil culturable microbes).

Principal coordinate analysis (PCoA) based on the Bray-Curtis distance matrix was used to evaluate the differences in microbial community structure among sugarcane. The analysis revealed a substantial resemblance in the endophytic culturable microbial communities within sugarcane, and there are no notable distinctions among sugarcane (Figure 2). This phenomenon indicates that the microbial communities within sugarcane plants are conserved, and may benefit both host and microbes through mutual interactions. On the other hand, there were significant differences in the composition of rhizosphere soil microbes and endophytic microbes in sugarcane. This can be attributed to the distinct ecological conditions between the rhizosphere soil and the interior of the sugarcane. The rhizosphere soil microbial community is more vulnerable to various factors, including soil fertility, environmental changes, and microbial interactions, in addition to the influence of the host plant. Furthermore, there were significant differences in the microbial communities in *S. robustum* and *S. officinarum* between different parts (Figs. 2B, 2C), reflecting that different parts of these two sugarcane species may be selectively enriched for specific microbial communities.

**Figure 2:**
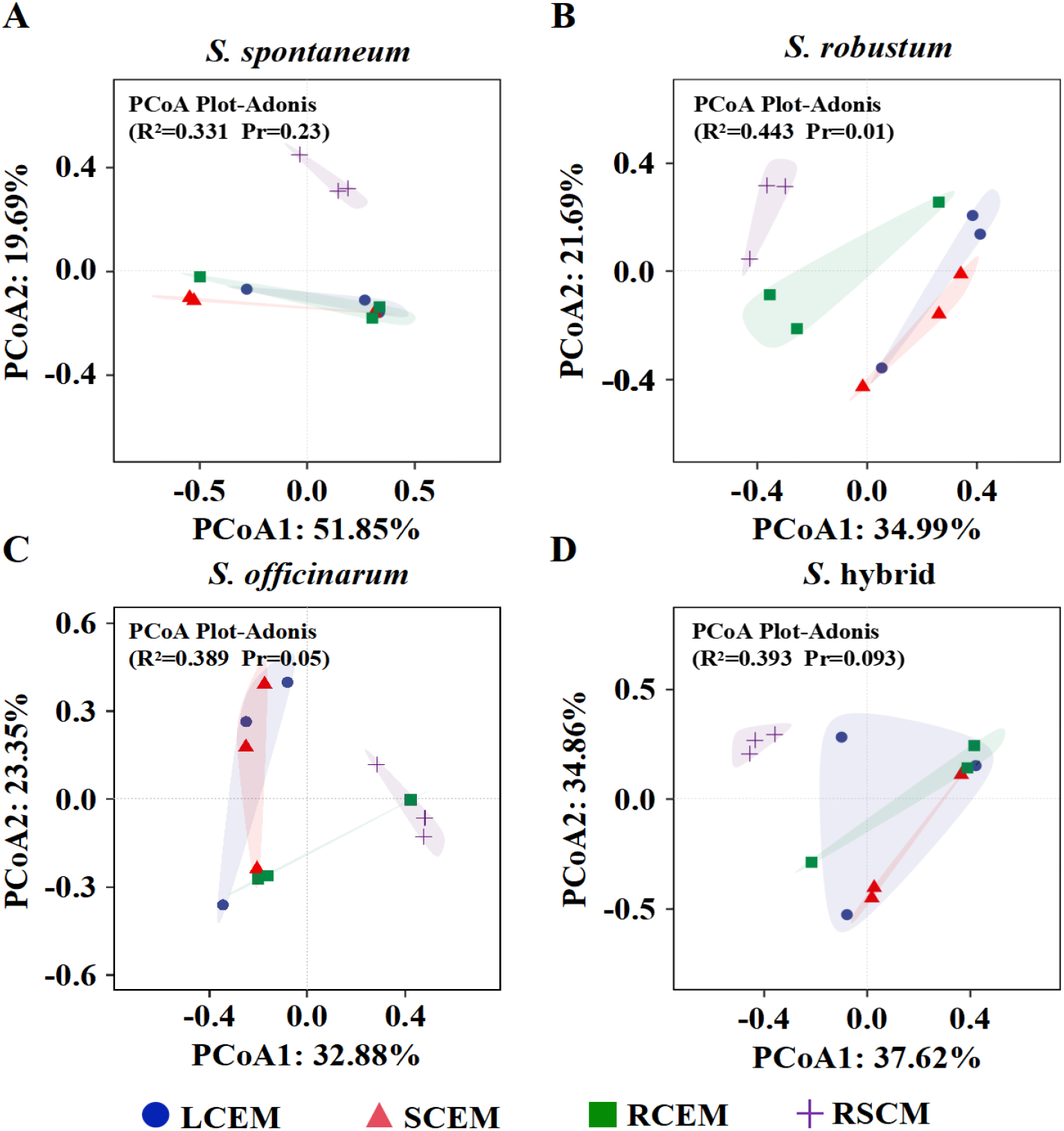
PCoA of culturable microbes from different compartments of sugarcane. R2 and P (Pr) values in the figure were based on Adonis calculations, **A, B, C**, and **D** indicated the differences in the structure of the culturable microbial communities among different compartments of *S. spontaneum, S. robustum, S. officinarum*, and *S*. hybrid.

### Analysis of the composition of culturable microbes in sugarcane

The investigation of endophytic and rhizosphere soil culturable microbes from sugarcane at the phylum level revealed the presence of 38 distinct bacterial phyla in these microbial communities. The dominant phyla in the culturable microbial community of sugarcane were *Pseudomonadota, Bacteroidota, Bacillota, Actinomycetota*, and *Cyanobacteria* (Figure 3A). These phyla collectively accounted for 99.9% of the community, indicating that they are highly adaptable and competitive in both sugarcane tissue and soil environments. In particular, *Pseudomonadota*, whose average relative abundance in the culturable microbial community of sugarcane was as high as 89.4%, was more distributed within the leaves than in other parts of the plant. Notably, the average relative abundance of *Pseudomonadota* isolated inside the leaves of the *S. robustum* and *S. officinarum* reached 97.9% and 98.1%, respectively, showing their dominance in the endophytic culturable microbial community of sugarcane, which may be related to their ability to rapidly adapt to the environment inside sugarcane leaves. In addition, *Bacteroidota* was also an important part of the culturable microbial phylum of sugarcane, with the highest relative abundance of 14.0% from the root of *S. robustum*.

**Figure 3:**
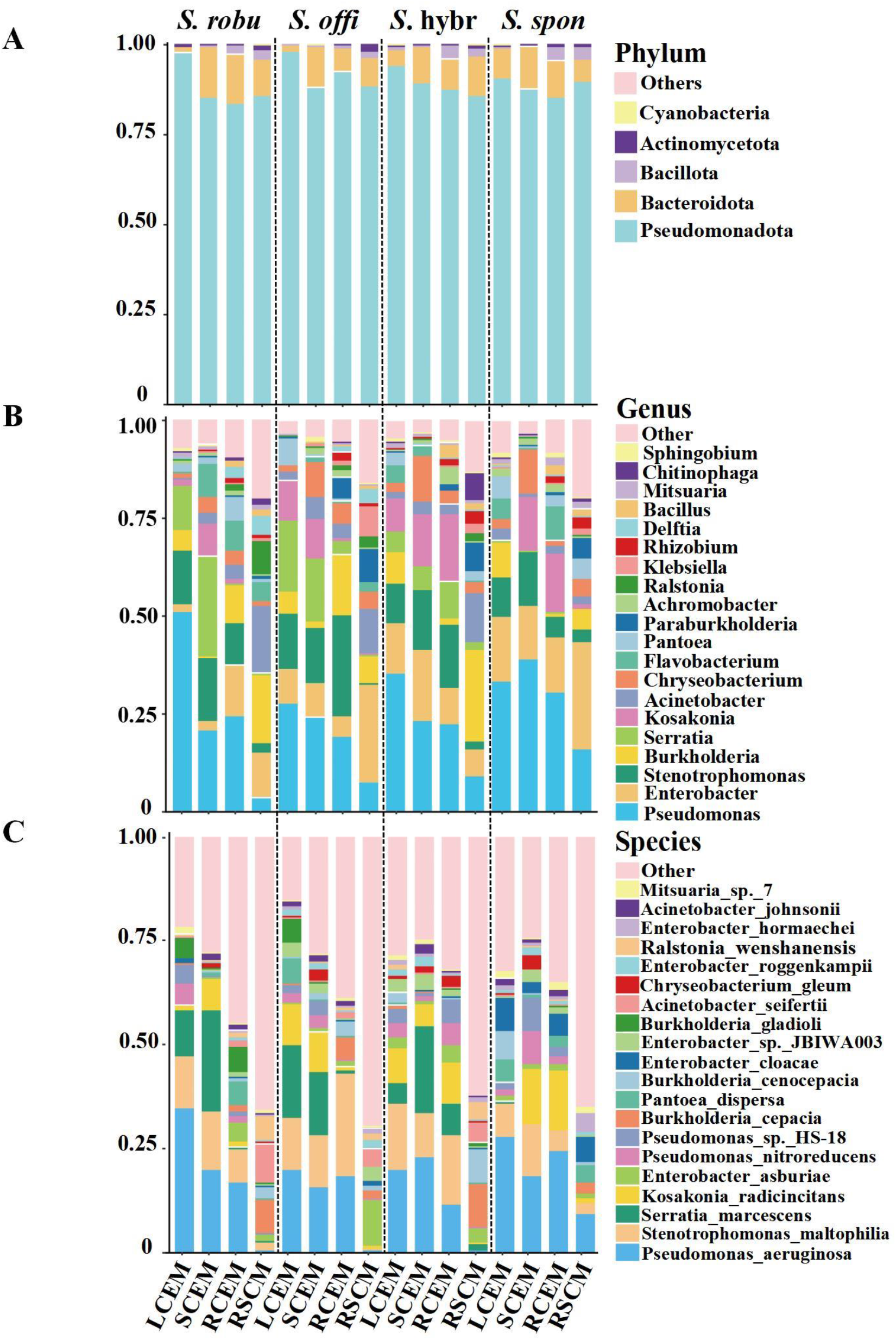
Composition of culturable microbes (phylum, genus and species level) from different compartments and sugarcane species.

Further microbial species annotation at the genus level revealed a total of 2274 genera identified, with the top five genera in terms of relative abundance under culture conditions including *Pseudomonas, Enterobacter, Stenotrophomonas, Burkholderia* and *Serratia* (Figure 3B). Microbes of these genera not only occupied an important position in the sugarcane microbial community but also showed distinct specific distribution patterns in different sugarcane species and their tissue sites. *Enterobacter* exhibited the highest relative abundance of 27.38% in the rhizosphere soil of *S. spontaneum*. In contrast, *Serratia* had the highest relative abundance of 25.5% within the stems of *S. robustum*, and *Burkholderia* had the highest relative abundance of 23.43% in the rhizosphere soil of the *S*. hybrid. *Pseudomonas* displayed a high relative abundance of 51.0% within the leaves of the *S. robustum* and an average relative abundance of 29.3% within the sugarcane. In contrast, this value was significantly reduced to 9.1% in the rhizosphere soil. This phenomenon was particularly significant in *S. spontaneum*, where the relative abundance of *Pseudomonas* in the rhizosphere soil was as high as 27.38%, significantly higher than that of other sugarcane species. This result suggests that *Pseudomonas* may have more suitable growth conditions in sugarcane or a closer mutualistic relationship with sugarcane, and also indicates that the root exudates or root structure of *S. spontaneum* is particularly attractive or supportive to *Pseudomonas*. In addition, the average relative abundance of *Stenotrophomonas* inside sugarcane was as high as 13.69%, in contrast to only 1.9% in the rhizosphere soil, which may imply that *Stenotrophomonas* are more adapted to grow in the internal environment of sugarcane.

Species annotation at the species level identified a total of 8271 species, and the results revealed *Pseudomonas aeruginosa, Stenotrophomonas maltophilia*, and *Serratia marcescens* to be the main dominant species (Figure 3C). The relative abundance of these dominant species in sugarcane showed differences among sugarcane species, such as *Burkholderia cepacia* and *Burkholderia cenocepacia* had the highest relative abundance in the rhizosphere soils of *S*. hybrid. *Serratia marcescens* had the highest relative abundance in the rhizosphere soil of *S. officinarum*, whereas it was almost undetected in all parts of *S. spontaneum*. In addition, *Pseudomonas aeruginosa* had the widest distribution, establishing its position as a dominant species with its average relative abundance of 16.2%. We were concerned that it presented a high abundance of 9.2% in the rhizosphere soils of *S. spontaneum*, whereas it was hardly detected in the rhizosphere soils of the other sugarcane. *Pseudomonas aeruginosa* was also abundant within the root system of *S. spontaneum*. The results suggest that *Pseudomonas aeruginosa* occupies a distinct ecological niche in the rhizosphere soil of *S. spontaneum*. Additionally, *S. spontaneum* may have a specific root-microbe mutualism that selectively recruits and enriches these beneficial microbes. This mutualistic relationship with *Pseudomonas aeruginosa* may contribute to the resilience and ecological competitiveness of *S. spontaneum* in the face of environmental stresses.

### Differences in the composition of culturable microbes in sugarcane

To examine the variations in the composition of microbes that can be grown in sugarcane, we conducted a comparative study of the microbes that can be cultured from different compartments (leaves, stems, roots, rhizosphere soil) of sugarcane (*S. spontaneum, S. robustum, S. officinarum*, and *S*. hybrid), respectively (Figure 4).The results showed that the *S*. hybrid species had the highest number of culturable microbial species shared between compartments, with a total of 5021 species (68.1%) (Figure 4D), suggesting that there was a high degree of consistency in microbial species of *S*. hybrid. In contrast, different compartments of *S. spontaneum* shared the lowest number of culturable microbial species, with a total of 4284 species (Figure 4A), which reflects that different parts of *S. spontaneum* may provide more diverse growth environments for microbes, thus promoting a more complex microbial community structure. The number of culturable microbial species unique to the rhizosphere soil of *S. spontaneum* reached 687 species (9.4%), which was significantly more than that of other sugarcane species, and this data emphasized the unique ecological characteristics of the rhizosphere soil environment of *S. spontaneum*. In *S. robustum*, the number of unique microbial species within the leaves was relatively low, only 37 species be founded, whereas the number of microbial species in the rhizosphere soil was as high as 326. For *S. officinarum*, fewer unique microbial species were found inside the leaves, only 25, compared to the rich microbial diversity in the rhizosphere soil, which was as high as 433 species. The lower microbial diversity in the endophytic environments of these sugarcane species compared to the rhizosphere soils may be related to the rich nutrients and environment required for microbial growth in the rhizosphere soils. Conversely, the endophytic environments exhibit fewer microbial species due to the constrained conditions they provide.

**Figure 4:**
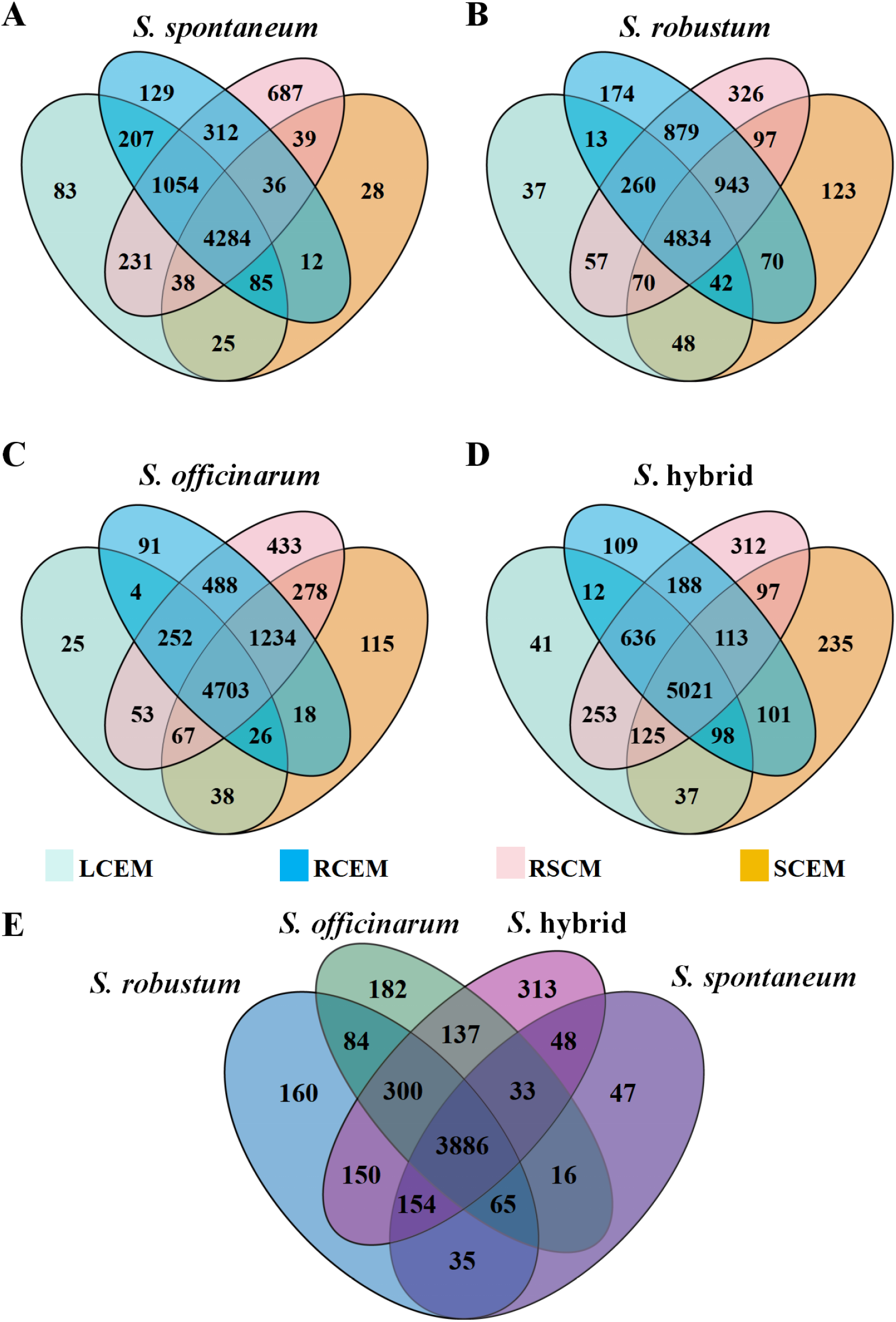
Unique and shared microbes from different compartments among sugarcane. **A, B, C** and **D** denoted the number of unique and shared culturable microbial species among different parts of *S. spontaneum, S. robustum, S. officinarum*, and *S*. hybrid, respectively. **E** denoted an ensemble of shared microbial species from sugarcane species

In a comparative analysis of the microbial communities of the four sugarcane species, we found a total of 3886 communal microbial species. In particular, *S. spontaneum* showed a significantly lower number of unique microbial species than the other species, with only 47 species. *S*. hybrid, which exhibited the highest number of unique microbial species, with 313 species (Figure 4E). This phenomenon indicates the richness of the *S*. hybrid species in microbial diversity. In contrast, *S. spontaneum* may have more excellent internal environmental stability and strict screening ability for microbes, and this unique microbial community structure may be related to its optimization of environmental adaptations and physiological functions, suggesting that *S. spontaneum* may have developed an efficient microbial mutualistic network.

LEfSe was used to analyze microbial differential markers in different parts of the same sugarcane species. The results demonstrated that *Burkholderia cenocepacia, Enterobacter sp JBIWA003*, and the rhizosphere soil of *Burkholderiaceae, Rhizobiacea*, and *Bacillaceae* exhibited LDA score values greater than 4 in different compartments of *S. spontaneum*, indicating that these microbes were notably abundant in *S. spontaneum* (Figure 5A). Similarly, microbial species with significant enrichment phenomena were found in different compartments of *S. robustum, S. officinarum* and *S*. hybrid (Figure 5B, C and D). Interestingly, the number of biomarkers that significantly differed within the stems of *S. officinarum* and within the leaves of *S*. hybrid relative to the other parts was 0, which may imply that the microbial community structure in these compartments is relatively stable. Notably, the LDA score values of *Betaproteobacteria, Burkholderiales, Burkholderiaceae* and *Burkholderia* in the rhizosphere soil of the *S*. hybrid were all greater than 5. This result suggests that these microbial taxa have significant relative abundance in specific compartments of *S*. hybrid. Furthermore, the *S. robustum* exhibited the most extensive range of biomarkers, with as many as 49 significantly different cultivable microbial markers. Additionally, it displayed LDA scores greater than 5 for *Yersiniaceae, Serratia* inside the stems, and *Burkholderiales* in the rhizosphere soil.

**Figure 5:**
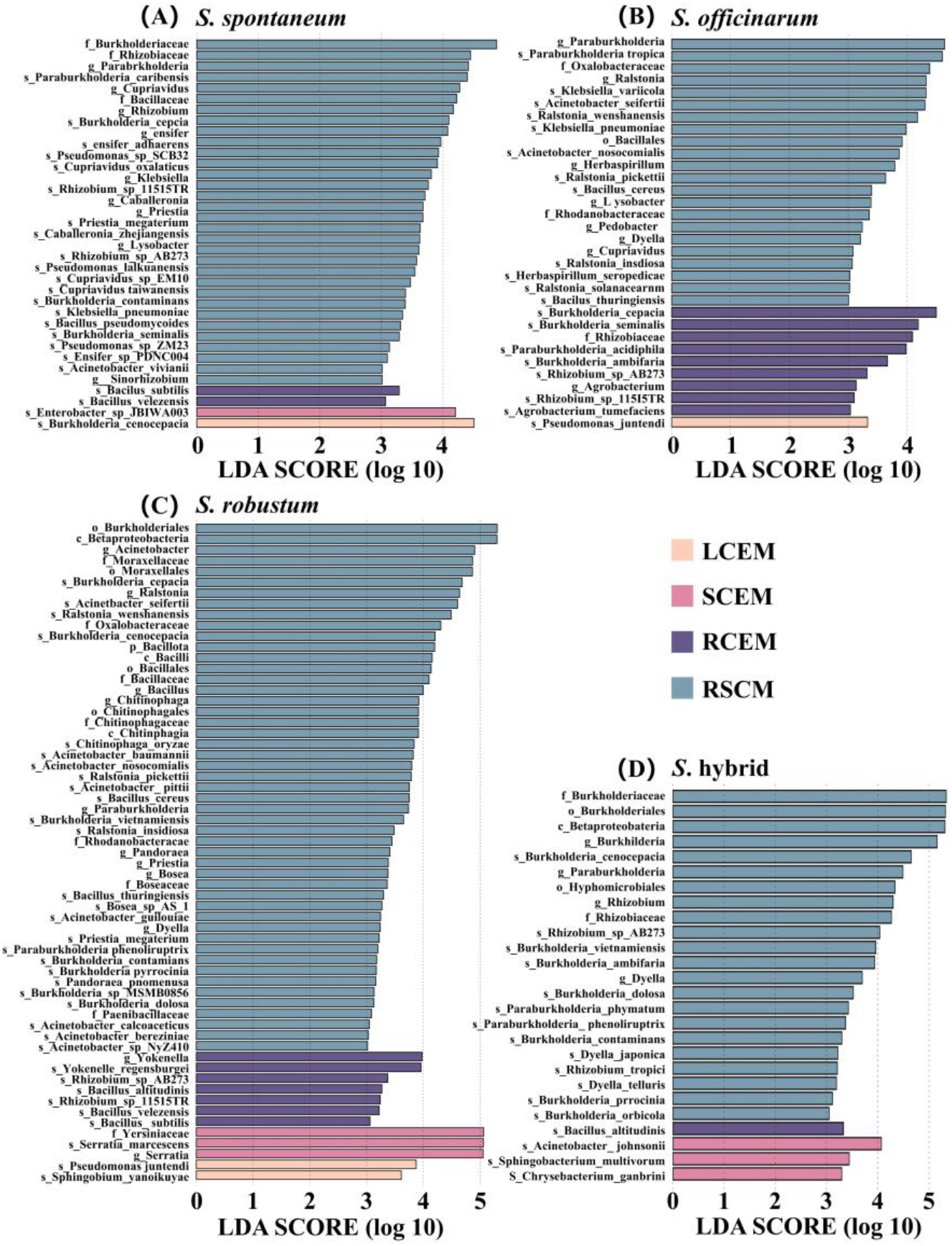
LDA Score of microbial significantly different species in different compartments of sugarcane. **A, B, C**, and **D** represented the distribution of LDA Score of culturable microbial significantly different species among different parts of *S. spontaneum, S. robustum, S. officinarum*, and *S*. hybrid, respectively.

### Culturable microbes in sugarcane may have the potential to regulate sucrose metabolism

Throughout the growth and metabolism of sugarcane, many tissues perform vital functions. They play a collaborative role in the production, transportation, storage, and metabolism of sucrose to fulfill the growth and developmental requirements of sugarcane. In order to gain a deeper understanding of how culturable microbes contribute to carbohydrate metabolism in different parts of sugarcane and their potential connections to sucrose metabolism, we performed comparative analyses of the functional sets of carbohydrase genes found in culturable microbial communities from sugarcane leaves, stems, roots, and rhizosphere soils. The PCoA analysis results further validated that the functional set of carbohydrase genes in the culturable microbes of sugarcane were grouped based on endophytes and rhizosphere soil microbes. Additionally, there was a notable impact of tissue parts on this clustering (PERMANOVA analysis: F=3.28, *P*<0.001) (Figure 6A), which aligned with the findings above based on the analysis of the community composition of culturable microbes in sugarcane.

**Figure 6:**
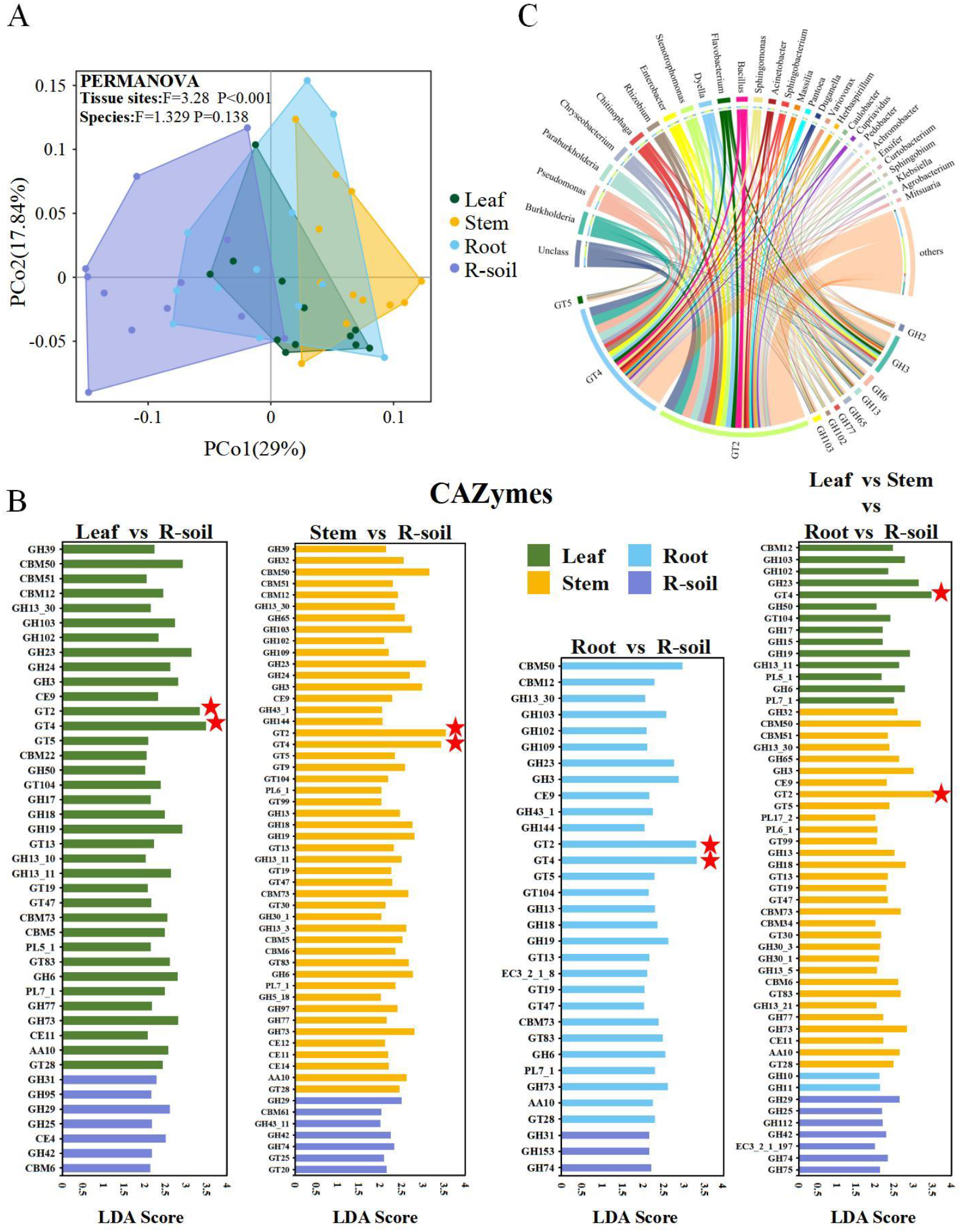
Functional differences of carbohydrases among different compartments of sugarcane. **A** Functional set of carbohydrase genes with significant abundance among different tissues of the genus sugarcane. Horizontal coordinates indicated LDA score values, vertical coordinates indicated different carbohydrases, and different colors represented tissues in the genus sugarcane. **B** Differences in carbohydrase gene function among culturable microbiota in different compartments of sugarcane. **C** CAZyes family with sucrose metabolism function (GT2, GT4, GT5, GH3, GH6, GH13, GH32, GH65, GH77, GH102 and GH103) associated with the top 20 microbial families with standardized relative abundance.

As revealed by LEfSe analysis, 36 (Leaf vs R-soil), 48 (Stem vs R-soil) and 29 (Root vs R-soil) carbohydrase markers were identified for culturable microbial carbohydrases within the tissues of sugarcane (Leaf, Stem, and Root) relative to rhizosphere soil culturable microbial carbohydrases, respectively (Figure 6B). Significantly, 23 carbohydrase markers exhibited a higher concentration in the cultivable microbes found in sugarcane tissues (leaves, stems, and roots). Among these markers, GT2 and GT4 had the highest level of enrichment (Figure 6B, Supplementary Table). The CAZy database showed that GT2 and GT4 were closely related to the sucrose metabolic pathway. Further analysis showed that these 23 carbohydrase markers, in addition to GT2 and GT4, GT5, GH3, GH6, GH32, GH65, GH77, GH102, and GH103, were all associated with the sucrose metabolic pathway. In particular, GT2 and GT4 showed higher enrichment levels in culturable microbes from the stem and leaf, respectively (Figure 6B). The analysis of the top 20 microbial families, in relation to their relative abundance compared to the CAZyme family showed that in the leaves and stems of sugarcane, genes for carbohydrate-activated enzymes (CAZymes), which are closely related to sucrose metabolism, were mainly attributed to several specific microbial genera, including *Burkholderia, Pseudomonas, ParaBurkholderia*, and *Chryseobacterium* (Figure 6C). These findings indirectly indicate that certain microbes and sugarcane have interactions. It suggests that these specific microbial groups may have important functions in sucrose metabolism in sugarcane.

## Discussion

In previous studies, researchers have typically used selective media to isolate single strains to study plant-associated microbes directly. Although this method is capable of isolating some beneficial bacteria, especially nitrogen-fixing bacteria (20), the number of strains isolated is small, limiting a comprehensive understanding of microbial diversity. Our research utilized metagenome sequencing to thoroughly evaluate the community composition of culturable endophytes from the root, stem, and leaf parts, as well as culturable microbes from the rhizosphere soil of four sugarcane species. This strategy avoids the issue of diminishing the diversity of cultivable microbes that might arise from conventional isolation methods.

This study found that different sugarcane germplasm and tissues were rich in endophytic culturable microbes. Specifically, there were differences in the diversity of culturable microbes in the same parts of sugarcane, but they were not statistically significant. This finding suggests that environmental conditions are as important as the characteristics of sugarcane itself when exploring differences in culturable microbial communities among sugarcane species (21, 22). Further PCoA showed similar endophytic culturable microbial community structures in different sugarcane germplasm from the same site. This suggests that in most cases, different germplasm of the same crop has relatively little effect on microbial community structure (23, 24). Among sugarcane’s endophyte compositions, *Pseudomonadota* dominated the sugarcane tissues. *Pseudomonadota* contains a wide range of bacteria with growth-promoting effects on plants, and they usually produce phytohormones, solubilise minerals, fix nitrogen and control plant diseases (25, 26). In the present study, we found that the average relative abundance of *Pseudomonadota* isolated from the leaves of *S. robustum* and *S. officinarum* reached 97.9% and 98.1%, respectively. In some studies, *Pseudomonadota* usually occupies the most dominant position of phyllosphere microbes (27, 28). This further highlights the centrality of *Pseudomonadota* in the endophytic microbial community of sugarcane leaves. In addition, this study observed specific significant differential enrichment of strains in sugarcane germplasm and tissue parts. For example, in sugarcane roots, significant differential enrichment of *Burkholderia vietnamiensis, Burkholderia ambifaria*, and *paraburkholderia tropica* was observed in *S. officinarum*, which all belong to the genus *Burkholderia. Burkholderia* is a genus of bacteria known to have several beneficial functions for plant growth such as nitrogen fixation, mineral solubilisation, synthesis of phytohormones and inhibition of plant diseases (29-31). This bacterium stimulates the production of biologically active substances, such as chitinase, cellulase, indoleacetic acid, and abscisic acid, in both the roots and above-ground parts of sugarcane (32). Inoculation with these bacteria regulates the metabolism of sugarcane and maintains its health (33). Specific dominant bacterial species in sugarcane tissues show significant survival advantages and form a mutually beneficial symbiotic relationship with sugarcane. Therefore, the study of strain differences between sugarcane species and parts provides important microbial resources for conducting thorough research on sugarcane’s interaction with its endophytes and lays the foundation for developing novel biofertilizers or biocontrol methods.

The results of PCoA analyses indicated the uniqueness of the rhizosphere soil culturable microbial community structure among sugarcane, which may be closely related to the specificity of different sugarcane germplasm and the composition of root exudates. Previous studies have shown that plant soil microbial community structure is influenced by factors such as plant root exudates, host plant and soil type (34-36). The community structure of sugarcane rhizosphere soil microbes was also influenced by sugarcane germplasm and its rhizosphere exudates (37). Interestingly, *S. spontaneum* showed greater differences in rhizosphere soil microbial community structure compared to the other species, whereas the soil-culturable microbial community structure among other sugarcane species had a high degree of similarity.

This may stem from the unique growth characteristics of *S. spontaneum* as a wild species of the genus sugarcane (*Saccharum* spp.). *S. spontaneum* are considered to be high-fiber plants with a significant geographic distribution that can survive a wide range of abiotic stress conditions such as drought, salinity, flooding and freezing (38). This study found that *Pseudomonas aeruginosa* is more widely present in the rhizosphere soil of *S. spontaneum* compared to other species. Additionally, *Pseudomonas aeruginosa* was highly abundant within the root system of *S. spontaneum*. These findings suggest that there may be a unique interaction mechanism between *Pseudomonas aeruginosa* and *S. spontaneum*’s root system. This mechanism could allow *S. spontaneum* to selectively recruit and enrich *Pseudomonas aeruginosa*, thereby enhancing its ability to adapt and compete in stressful environmental conditions.

In this study, we identified significantly enriched bacterial markers in sugarcane by LEfSe analysis, particularly *Paraburkholderia caribensis* and *Enterobacter bugandensis*, which were significantly enriched in rhizosphere soils of *S. spontaneum*. These bacteria have been reported in other plant systems to have nitrogen-fixing capacity and enhance plant tolerance under drought conditions (39-41). In the rhizosphere soil of *S. spontaneum*, a large number of bacterial genera with nitrogen-fixing capacity and disease suppression have also been discovered (42). Significant enrichment of the bacterial marker *Burkholderia* was identified in the roots of *S. officinarum* relative to other sugarcane species. Also, multiple significantly enriched strains of *Paraburkholderia* and *Burkholderia* were present in roots and rhizosphere soil of *S. officinarum* relative to leaves and stems. *Paraburkholderia* and *Burkholderia* are highly versatile bacteria that inhabit a wide range of ecological niches in various hosts. They are known for their exceptional ability to enhance plant growth, particularly in the rhizosphere. These bacteria are commonly found in significant quantities in major crops such as maize, rice, wheat, tomato, and potato, contributing to plant growth through processes such as biogenic nitrogen fixation, production of iron carriers, solubilization of inorganic phosphate, production of indole acetic acid, and inhibition of plant pathogens (43-45). Notably, the roots and rhizosphere soil of *S*. hybrid contained a significant amount of *Paraburkholderia* and *Burkholderia* strains. The *S*. hybrid is derived from interspecific crosses between *S. officinarum* and *S. spontaneum* and backcrosses with *S. officinarum* backcrosses, which contribute to increased sugar content and stress tolerance (46-48). This provides more evidence that *S*. hybrid continues to possess advantageous characteristics inherited from its parents, which have significant and widespread impacts on the growth and development of sugarcane. The results demonstrated the presence of distinct microbial enrichment in various parts of different sugarcane species. This finding highlights the complex interactions between sugarcane and microbes, as well as the diversity and specificity of sugarcane microbial communities. Therefore, mining sugarcane-specific microbes holds great importance for the domestication, breeding, and cultivation of sugarcane.

There were notable variations in the types of carbohydrate enzymes found in the culturable microbes present in sugarcane tissues. Sucrose, the primary sugar in sugarcane, is produced in the leaves and transported to the stem. Initially, there is a small amount of sucrose in the upper immature internodes, but as the sugarcane matures, it progressively accumulates in the lower internodes. Eventually, sucrose can make up around 50% of the total dry weight of the sugarcane (49). Previous studies have shown that nitrogen-fixing bacteria isolated from sugarcane roots or soil prefer to grow in media with high sucrose concentrations (50). In this study, we found that the relative abundance of *Burkholderia* with nitrogen-fixing ability was 4.95% in the tissues. Thus, the current study determined that sugarcane tissues with a high sucrose content provide an optimal habitat for the colonization of particular endophytes. Out of all the bacteria that can be cultivated and have the ability to metabolise sucrose in sugarcane, *Burkholderia* was found to be the most prevalent group, suggesting that *Burkholderia* may have a connection with the accumulation and metabolism of sugar in sugarcane. Sugarcane tissues exhibiting high sugar content have a strong propensity to attract a substantial population of *Burkholderia* for colonisation. Sugarcane provides a rich source of sucrose carbon for *Burkholderia* bacteria to support their growth, development and reproduction. On the other hand, *Burkholderia* bacteria may promote nutrient utilisation and growth of sugarcane, thereby increasing sucrose production from sugarcane and creating a mutually beneficial relationship. Additional analyses of the connections between high-sugar sugarcane and the microbiome will contribute to establishing a scientific foundation and theoretical direction for utilising microbial fertilisers to enhance the sucrose production of sugarcane, and advance the sustainable growth of the sugarcane industry.

